# Graded spikes differentially signal neurotransmitter input in cerebrospinal fluid contacting neurons of the mouse spinal cord

**DOI:** 10.1101/2020.09.18.303347

**Authors:** Emily Johnson, Marilyn Clark, Merve Oncul, Claudia MacLean, Jim Deuchars, Susan A. Deuchars, Jamie Johnston

## Abstract

The action potential and its all-or-none nature is fundamental to neural communication. Canonically the action potential is initiated once voltage-gated Na^+^ channels are activated, and their rapid kinetics of activation and inactivation give rise to the action potential’s all-or-none nature. Here we show that cerebrospinal fluid contacting neurons (CSFcNs) surrounding the central canal of the mouse spinal cord employ a different strategy. Rather than using voltage-gated Na^+^ channels to generate binary spikes, CSFcNs use two different types of voltage-gated Ca^2+^ channel, enabling spikes of different amplitude. T-type Ca^2+^ channels generate small amplitude spikes, whereas large amplitude spikes require high voltage-activated Cd^2+^ sensitive Ca^2+^ channels. We show that these different amplitude spikes signal input from different transmitter systems; purinergic inputs evoke smaller T-type dependent spikes while cholinergic inputs evoke large T-type independent spikes. Different synaptic inputs to CSFcNs can therefore be signalled by the spike amplitude.

## Introduction

Cerebrospinal fluid contacting neurons (CSFcNs) surround the central canal of the spinal cord in all vertebrate species examined (Djenoune et al., 2014) and possibly in humans (Humphrey, 1947). They project a single dendritic like structure into the CSF through the ependymal cells that form the border of the central canal. CSFcNs are also present in the caudal medulla oblongata, predominantly surrounding the central canal (Orts-Del’Immagine et al., 2017; Orts-Del’immagine et al., 2012a).

Upon their identification it was suggested that CSFcNs form a sensory “sagittal organ” within the spinal cord (Kolmer, 1921). This idea is consistent with the observation that CSFcN are the only cells in the CNS that express polycystic kidney disease 2-like 1 protein (PKD2L1), a channel reported to have chemo- and mechanosensitive properties (Huang et al., 2006; Djenoune et al., 2014; Orts-Del’Immagine et al., 2012b; Orts-Del’Immagine et al., 2014; Orts-Del’Immagine et al., 2016). Indeed, studies in zebrafish have indicated that different CSFcN populations respond to bending of the spinal cord and synapse onto distinct motor neuron populations and afferent interneurons to regulate motor behaviour. In these studies, disruption of CSFcN signalling leads to impairment of postural control (Hubbard et al., 2016), spinal morphogenesis (Sternberg et al., 2018) and locomotion (Wyart et al., 2009; Fidelin et al., 2015; Böhm et al., 2016; Hubbard et al., 2016). Recordings from lamprey also indicate that CSFcNs play a homeostatic role in locomotion; CSFcNs were sensitive to pH and deviations from normal pH reduced locomotor output (Jalalvand et al., 2016b; Jalalvand et al., 2016a). These findings indicate a key role for CSFcNs in spinal sensory signalling in lower vertebrates, yet little is known about the function or indeed the signalling mechanisms of these cells in mammalian systems.

Voltage-activated Na^+^ channels are widely considered to be fundamental for neuronal excitability and are a requirement for the generation and propagation of action potentials throughout the central and peripheral nervous systems (Wang et al., 2017). Whilst this assumption holds true in most mammalian neurons, sensory systems commonly utilise Ca^2+^ as the primary mediator of electrogenesis. Within auditory hair cells Ca_V_1.3 (L-type) channels can mediate spikes and glutamate release (Brandt et al., 2003; Zampini et al., 2010). Similarly, retinal bipolar cells rely on low-voltage activated (T-type) Ca^2+^ channels to initiate regenerative potentials and spiking activity (Hu et al., 2009; Dreosti et al., 2011). As voltage activated Ca^2+^ channels operate over a wide range of membrane potentials and facilitate both spiking and graded events, their prevalence enables sensory neurons to respond to a wide range of inputs (Lipin and Vigh, 2015; Baden et al., 2013). Such a mechanism for signalling would also be advantageous for CSFcNs if they fulfil a sensory role. Single-cell RNA sequencing indicates that mouse spinal CSFcNs abundantly express mRNA for all 3 isoforms of T-type calcium channel (Ca_V_3-1, -2, -3) and numerous high-voltage-activated (HVA) Ca^2+^ channels, including Ca_V_1.3 (Rosenberg et al., 2018) an L-type channel predominantly expressed by sensory neurons and neurosecretory cells (Comunanza et al., 2010; Joiner and Lee, 2015).

To begin addressing whether CSFcNs constitute a novel sensory system within the mammalian spinal cord we used 2-photon Ca^2+^ imaging to study the activity of mouse CSFcNs. Our findings reveal that mouse spinal CSFcNs exhibit T-type Ca^2+^channel dependent spontaneous activity and, in parallel to other sensory systems, employ voltage-activated Ca^2+^ channels to generate spikes of graded amplitudes. CSFcNs can use this amplitude code to signal which of their neurotransmitter systems have been activated.

## Results

### The VGAT promotor drives GCaMP6f expression in all CSFcNs

To enable imaging of neural activity within CSFcN populations, we targeted GCaMP6f to CSFcNs of the central canal by driving its expression under the VGAT promoter. Within the spinal central canal GCaMP6f was expressed by cells with stereotypical CSFcN morphology, displaying a single bulbous apical process extending into the lumen of the central canal (Fig 1). PKD2L1, the canonical marker of CSFcNs (Djenoune et al., 2014), displayed 100% overlap with these GCaMP6f positive cells (Fig. 1, n= 206, N=6) and all VGAT+ cells with stereotypical CSFcN morphology were positive for PKD2L1. These data indicate that GCaMP6f is expressed in the entire population of CSFcNs in our VGAT-GCaMP6f mice, we next took advantage of 2-photon microscopy to image their activity in acute spinal cord slices.

**Figure 1:**
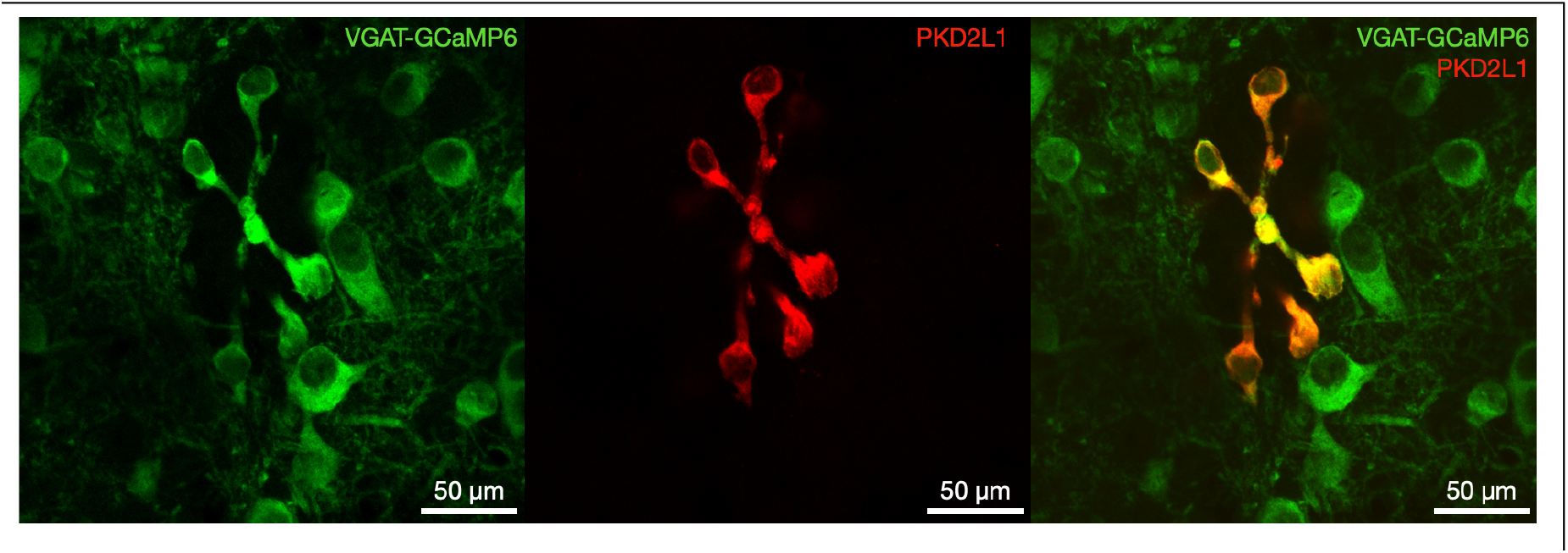
All PKD2L1 expressing CSFcNs are GABAergic. *left* VGAT-GCaMP6 expression around the central canal of the spinal cord. *Middle* PKD2L1 expression in the same section. *Right* composite image showing 100% overlap between PKD2L1 and VGAT-GCaMP6. Doral side is at the top. Representative data from 6 animals, 12 sections per animal.

### CSFcNs generate variable amplitude spikes

We observed widespread spontaneous activity across the population of CSFcNs (Fig 2A, B) similar to that reported from *in vivo* Ca^2+^ imaging in the larval zebrafish (Sternberg et al., 2018). We detected Ca^2+^ spikes using their first derivative (Fig. 2C) which enabled separation of summated spikes, see methods for further details. Spontaneous activity occurred at a low frequency in CSFcNs (Fig. 2B & G); across a population of 127 CSFcNs the firing rate was 0.148 Hz (median, IQR = 0.097 Hz, n=127, N=15). The coefficient of variation for the inter-spike-interval of CSFcNs was close to 1, indicating that their spontaneous activity can be described by a simple Poisson process (Fig. 2H). Strikingly, individual CSFcNs displayed spikes of variable amplitude with distinct low and high amplitude events detected (Fig. 2 B, D & E), 81 of the 127 CSFcNs recorded showed such multimodality (Fig. F). These properties appeared stable with age as neither the spike rate, CV of the inter-spike interval nor the multimodality were correlated with age across the range we tested, P19 to 47 (p = 0.54, p = 0.21 and p = 0.38 respectively, supplementary figure S1). The larger Ca^2+^ spikes we observe may simply result from multi-action potential events; the expectation would then be that larger events would have slower rise times, yet despite large variation in amplitude the spike waveform was remarkably consistent (Fig. 2D). This led us to explore the electrical activity underlying these spikes.

**Figure 2:**
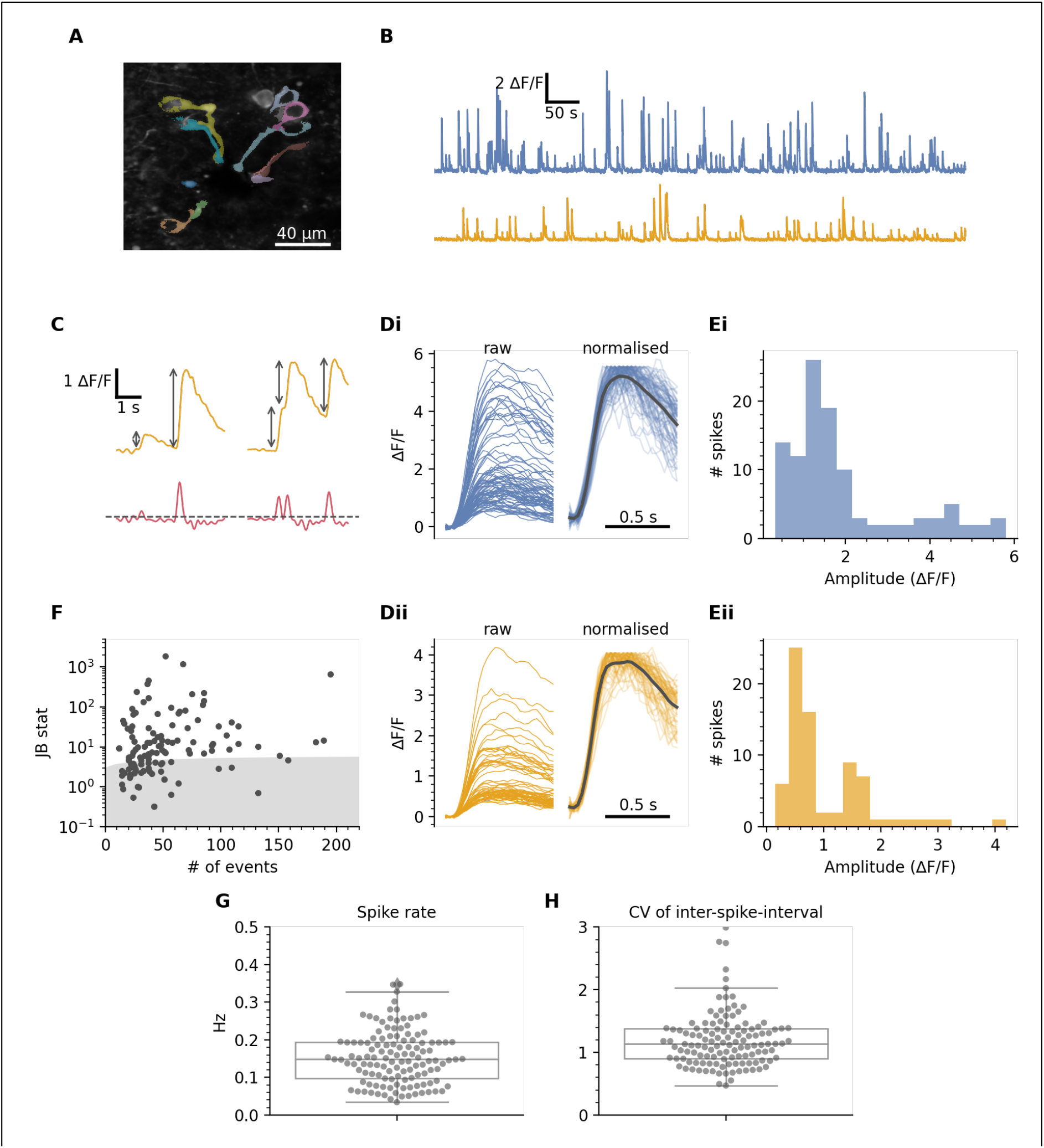
CSFcN spontaneous activity. **A)** Field of view containing CSFcNs showing segmentation of different CSFcNs. **B** Spontaneous activity of 2 example cells from A. **C** Spikes were detected on the differentiated trace (red) with an automatically determined threshold (see methods), grey arrows indicate amplitude measurements. **D** The parsed spikes from B, aligned to their onset, the left shows the raw ΔF/F signal and the right shows the spikes normalised to their max. Black line shows the mean signal for normalised spikes. **E** Amplitude histograms for the data in D. **F** The spike amplitudes within each cell was not normally distributed in 81 of 127 cells, grey shaded area shows critical value of the Jarque-Bera (JB) statistic, see methods. **G** Spontaneous rates from 127 CSFcNs from 15 animals had a median of 0.148 ±0.09 (IQR) **H** The coefficient of variation of the inter-spike intervals across the population was 1.13 ± 0.48 (median ±IQR, n=127, N=15).

### Variable recruitment of HVA Cav channels generates variable amplitude spikes

To investigate the source of the amplitude variability in these Ca^2+^ spikes we began by recording extracellular action potentials (EAPs) from identified CSFcNs. EAPs measure the membrane currents associated with the action potential (Rall, 1962; Holt and Koch, 1999; Gold et al., 2006; Perkins, 2006) and are a close approximation to the 1^st^ derivative of the intracellular action potential (Matthews and Lee, 1991; Buzsáki et al., 1996). Figure 3A illustrates a typical EAP waveform and its integral recorded from a non-CSF contacting neuron (black) and from a CSFcN (red). CSFcN action potentials had slow rise times of 3.9 ± 1.5 ms (n=16, N=10, mean ± SD), consistent with previous recordings (Corns et al., 2015; Marichal et al., 2009). Strikingly, CSFCNs displayed a dual peak in the depolarising phase of their EAP, with the secondary peak having an amplitude 17 ±3% of the primary peak (calculated from the mean spike waveforms, n = 16, N=10). This secondary peak also showed the highest spike-to-spike variability (Fig. 3B & C); it wasn’t evident with every spike and could be as large as the initial peak (Fig. 3B bottom). Similar secondary depolarising peaks are observed in the EAPs of other slow spiking cells and are due to activation of Cd^2+^ sensitive HVA Ca^2+^ channels (Matthews and Lee, 1991). The variable secondary depolarising phase we observe may therefore be due to differential recruitment of HVA Ca^2+^ channels, a potential source of the amplitude variability of Ca^2+^ spikes. Consistent with this idea 100 µM Cd^2+^, a broad-spectrum HVA Ca^2+^ channel blocker, caused a hyperpolarising shift in the secondary peak of the EAP in CSFcNs by 5.9 ±2.7 pA (mean ±SD, n=4, N=4, Fig. 4 A & B). Correspondingly, Cd^2+^ caused a significant reduction in the mean amplitude of spontaneous Ca^2+^ spikes (Control: 1.54 ΔF/F IQR = 3.79 vs +Cd^2+^: 1.04 ΔF/F IQR = 1.39, p = 1.18×10^−6^, n = 65, N = 5, Figs. 4 C-E). This reduction in the mean amplitude was a result of the larger amplitude spikes being inhibited which can be seen in the single example in shown in Fig. 4C-D and in the normalised amplitude histograms for all spikes from 65 cells (Fig. 4F). Together these data indicate that the variable amplitude Ca^2+^ spikes shown in figure 2 are underpinned by variable recruitment of HVA Ca^2+^ (Fig. 3 & 4).

**Figure 3.**
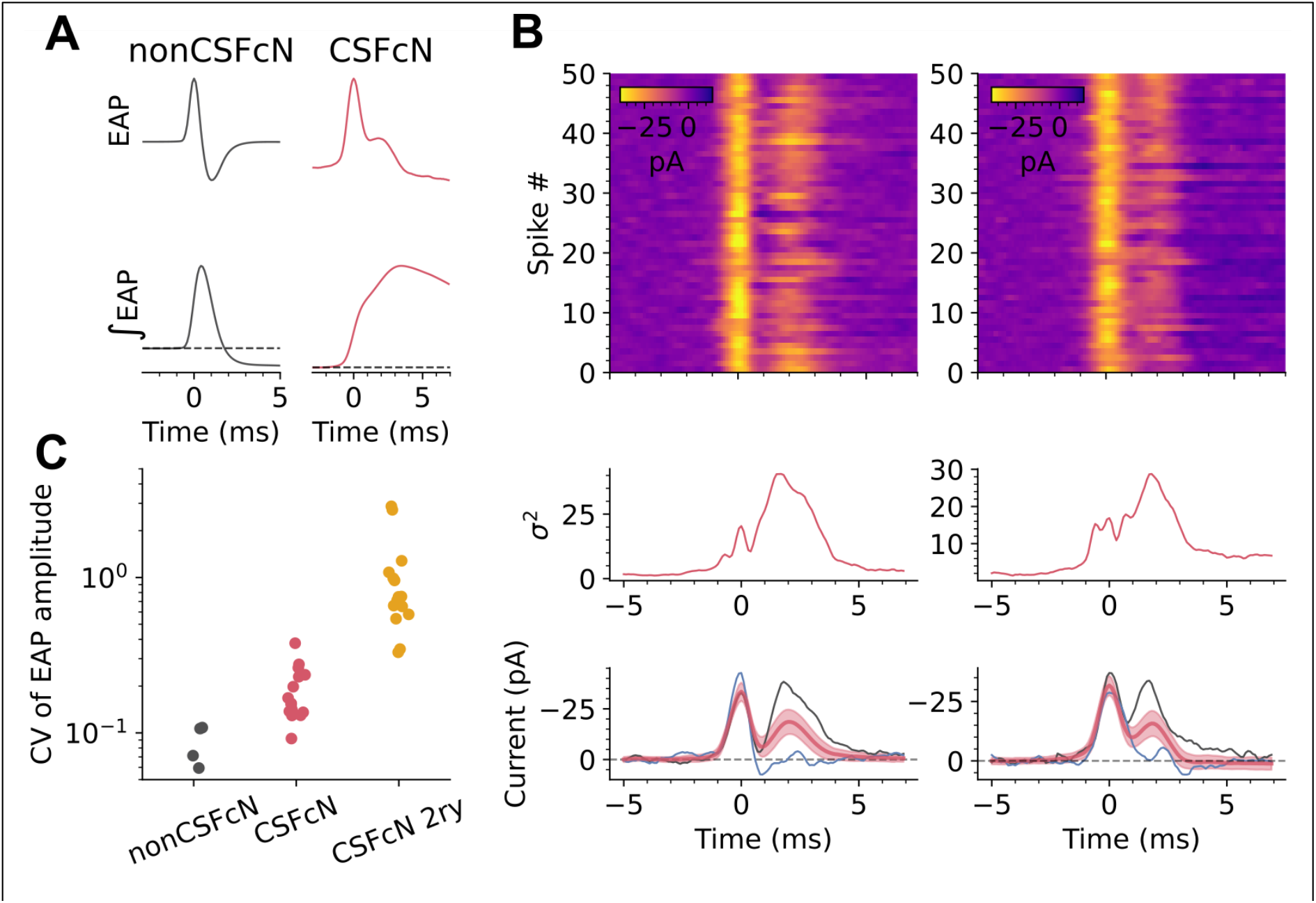
CSFcNs display variable action potential waveforms with a dual depolarising phase. **A)** Example of an EAP (top row) recorded from a non-CSFcN (black) and a CSFcN (red) and their integrals (bottom row), CSFcNs in EAP typically display a secondary depolarising peak. **B)** Raster plots showing 50 EAPs from two further CSFcNs (top row) and the variance of the waveforms (middle row) with the mean and standard deviation (red traces, bottom row), with individual examples in black and blue. The secondary depolarising peak showed the largest variance and could be as large as the initial peak (black examples, bottom row) or almost absent (blue traces, bottom row). **C)** The coefficient of variation in the amplitude of the depolarising peaks of the EAP from non-CSFcNs (n=4,N=4) and CSFcNs (n=16, N=16).

**Figure 4.**
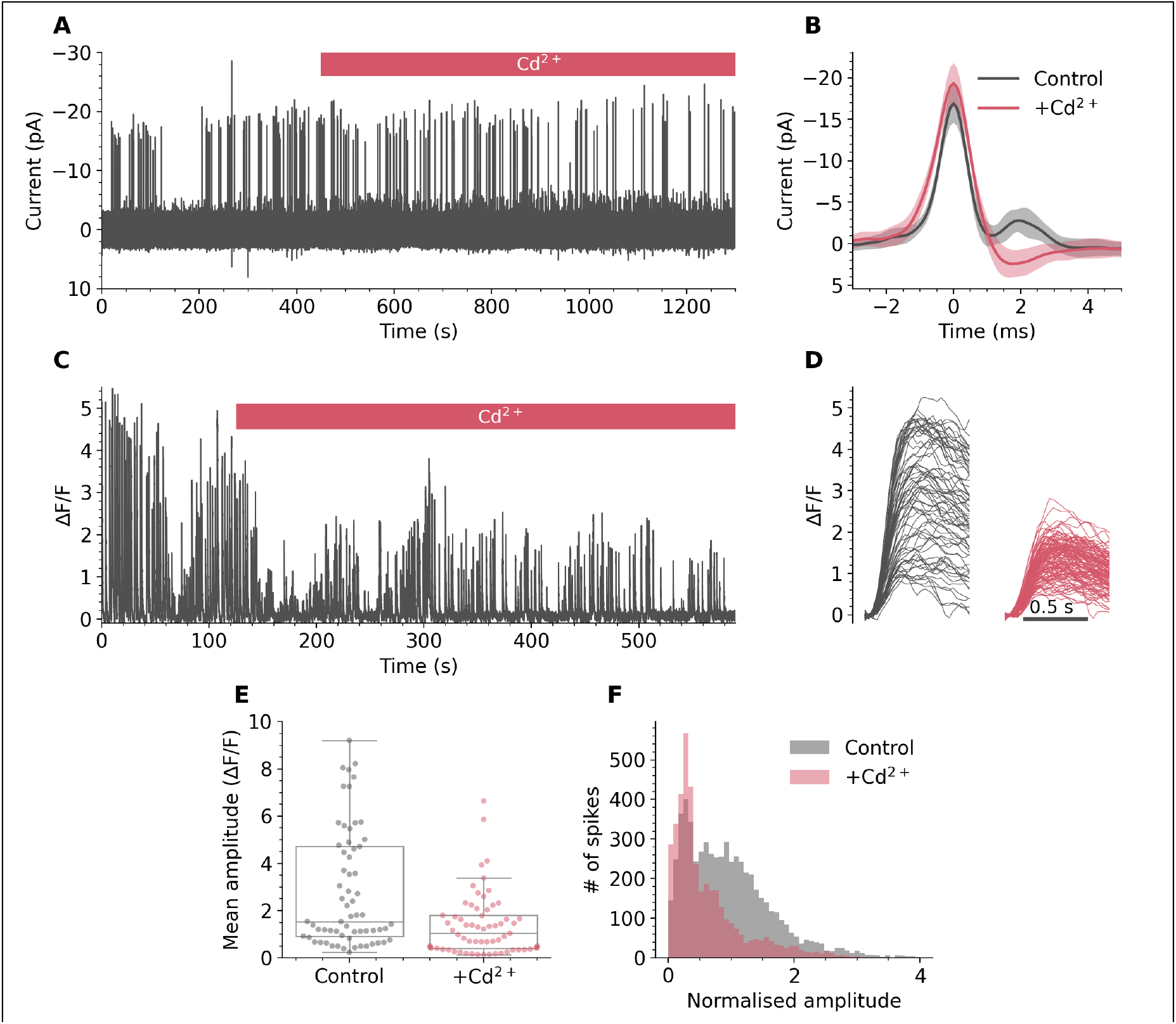
High-voltage activated Ca^2+^ channels generate the secondary depolarising peak in CSFcN EAPs and large amplitude Ca^2+^ spikes. **A)** EAP recording from a CSFcN during application of 100 µM Cd^2+^. **B)** The mean ± standard deviation of EAP wave form of the first 50 events prior to Cd^2+^ application (black) and the last 50 events in the presence of Cd^2+^ (red). Cd^2+^ caused a hyperpolarsing shift in the secondary peak of 5.4 ±1.9 pA (n=4, N=4). **C)** Ca^2+^ spikes recorded from a CSFcN during application of 100 µM Cd^2+^. **D)** The Ca^2+^ spikes from C parsed out before (black) and after (red) Cd^2+^ application. Note the variable amplitude in control (black) which become smaller and less variable in Cd^2+^. **E)** 100 µM Cd^2+^ significantly reduced the mean amplitude of CSFcN Ca^2+^ spikes (n=65, N=5, p = 1.18×10^−6^, Wilcoxon signed rank). **F)** Histograms of CSFcN amplitudes normalised to the mean amplitude in control. A significant reduction in large amplitude events occurs with Cd^2+^ (n=65, N=5, p = 9.33 x10^−15^, Kolmogorov-smirnov test).

### Mechanical activation of CSFcNs by the recording electrode

We next sought to correlate the EAP waveform with simultaneously measured Ca^2+^ spikes in CSFcNs. Although the cell-attached recording configuration is minimally invasive, with the intracellular milieu remaining completely unaltered (Perkins, 2006), we found the mean spike rate to be 3-fold higher when measured with EAPs compared to Ca^2+^ imaging (EAP rate 0.43 ±0.18 Hz, mean ±SD, n=15, N=10, vs Ca^2^ spike rate: median 0.148 ±0.097 Hz median ±IQR, n = 127, N=15). Strikingly the different rates we measured with EAPs and Ca^2+^ imaging are almost identical to previous electrical recordings comparing spontaneous activity in wildtype (0.42 Hz) vs animals lacking the mechano-sensitive PKD2L1 channel (0.16 Hz) (Orts-Del’Immagine et al., 2016). This indicates that the higher firing rate we observe with EAPs may be a result of the mechanical perturbation resulting from the necessity of placing a patch electrode against the cell. Consistent with this idea, only the patched cell showed elevated activity. Figure 5 shows the activity of 4 CSFCNs in a field of view, pre and post placement of a patch electrode to record EAPs from the cell in blue. Initially the blue cell is spiking at a typical low rate, once the patch electrode is in place the basal Ca^2+^ remains elevated and the measured EAP rate was 0.92 Hz, despite great care in preventing any depolarisation via the pipette (Perkins, 2006); the other cells in the field of view were unaffected. Higher levels of Ca^2+^ activity were observed with imaging in all 15 cells with simultaneous EAP recordings with average activity of 1.43 ± 1.01 ΔF/F s^-1^ (n=15) for EAP cells compare to 0.13 ± 0.7 ΔF/F s^-1^ (n= 66, Fig. 5 C) for their neighbours. In summary, placement of an electrode against CSFcNs elevates their spike rate and Ca^2+^ activity, this is likely due to the mechano-sensitive, large conductance and Ca^2+^ permeable PKD2L1 channels expressed in these cells (Fig. 1) (Böhm et al., 2016; Sternberg et al., 2018). As the majority of cells had aberrant Ca^2+^ activity it precluded us from systematically comparing electrical spikes with Ca^2+^ spikes. Figure 6 shows an example of one of the five cells where we were able to directly compare the EAP and Ca^2+^ spikes due to the Ca^2+^ activity being less affected by the EAP electrode. All fast-rising Ca^2+^ spikes detected had corresponding EAPs (Fig. 6A red dots Ca^2+^ spike detection). Equally all EAPs were accompanied by a fast-rising Ca^2+^ spike (Fig. 6A, yellow dots EAP detection). Interestingly a few small amplitude and slow rising Ca^2+^ events (red arrows Fig. 6A) were not accompanied by EAPs. Due to their slow rate of rise these events were not picked up by our spike detection criteria, they may reflect slower electrical events or be a result of spontaneous PKD2L1 openings that are observed when an electrode is in place on a CSFcN (Orts-Del’Immagine et al., 2016). Figure 6 B shows the Ca^2+^ spikes expanded with their corresponding EAPs shown below, with a further expanded time base. Similar to figure 3, the presence of the secondary depolarising peak of the EAP varies. The peak of the EAP integral, which is proportional to the action potential amplitude (Matthews and Lee, 1991; Buzsáki et al., 1996), was correlated with the amplitude of the Ca^2+^ spike (Fig. 6C), the R^2^ for this cell was 0.76 and all 5 cells showed a significant correlation with a mean R^2^ of 0.51 ±0.14 (Fig. 6D). This shows that a large fraction of the variance in Ca^2+^ spike amplitude is accounted for by differences in the underlying action potential. A significant proportion of the remaining variance is likely explained by noise as, across the 5 cells, the R^2^ value for EAP vs Ca^2+^ spike was correlated with the signal-to-noise ratio of the Ca^2+^ spikes (R^2^ = 0.64, Fig. 6D). Similar to figure 2D, the Ca^2+^ spikes had a stereotypical rising phase, despite over a 4-fold difference in amplitude, which can be seen in figure 6E where we normalised the Ca^2+^ spikes to their amplitude 200 ms after the EAP; the GCaMP6f sensor has a time-to peak of 200 ms (Chen et al., 2013). Together, these data show that placement of the patch electrode against CSFcNs results in higher basal electrical and Ca^2+^ activity (Fig. 5) consistent with previous reports of mechanosensation in CSFcN of zebrafish and lamprey (Sternberg et al., 2018; Jalalvand et al., 2018). These data also show that the Ca^2+^ spikes characterised in figure 2 are due to underlying electrical spikes and their amplitude variability is correlated with the presence of a secondary depolarising peak in their action potential (Fig. 6) which is mediated by HVA Cd^2+^ sensitive Ca^2+^ channels (Fig. 4).

**Figure 5:**
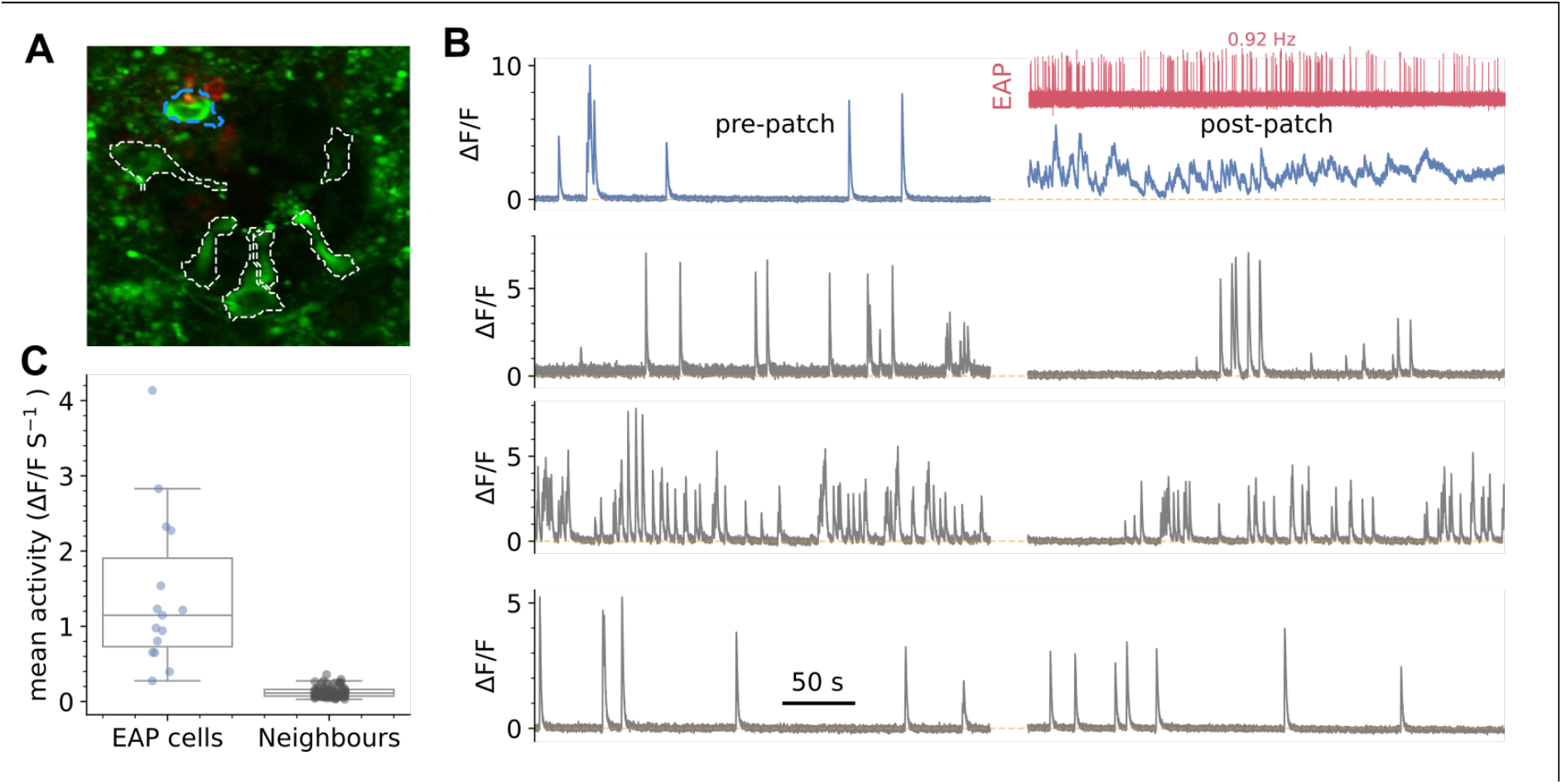
Mechanical activation of CSFcNs via the patch pipette. **A)** VGAT-GCaMP6 image of CSFcNs (green and dashed outlines) with an EAP electrode against the blue dashed cell. **B)** top: spontaneous activity of the blue cell in A before and after placement of the EAP electrode, red trace shows the EAP activity. Bottom grey rows show 3 neighbouring CSFcNs recorded simultaneously in the same field of view that didn’t have an electrode placed against them. **C)** Ca^2+^ activity was higher in all cells with a EAP electrode compare to their neighbours EAP cells (1.43 ± 1.01 vs 0.13 ±0.7 ΔF/F s^-1^, p = 1.4×10^−9^, Mann-Whitney-U test, EAP n = 15, neighbours n = 66)

**Figure 6:**
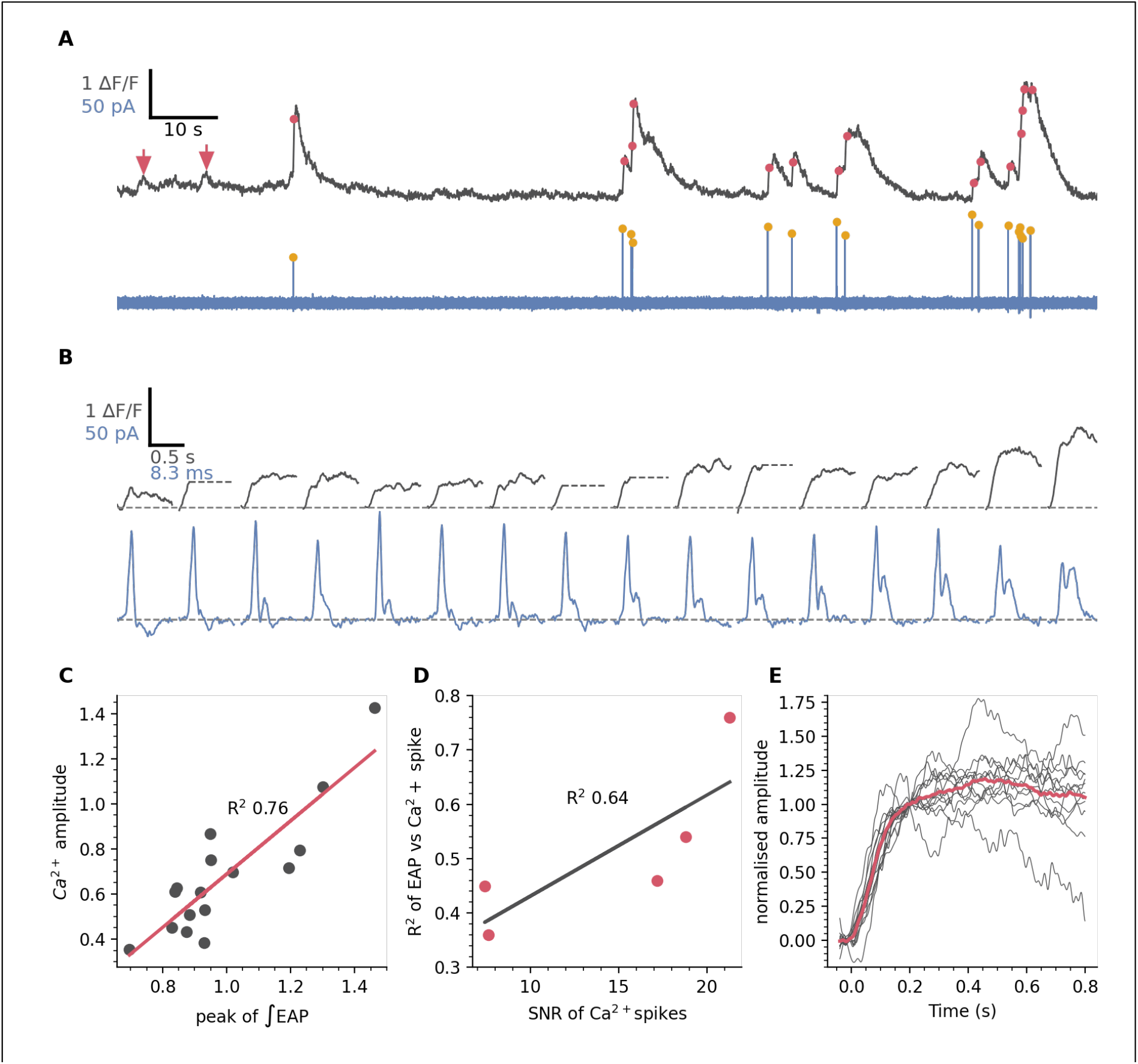
EAP waveform is correlated with Ca^2+^ spike amplitude. **A)** Simultaneously recording of Ca^2+^ activity (grey) and EAPs (blue). Detected EAPs indicated by yellow dots and Ca^2+^ spikes detected as in figure 2 are indicated by red dots. Note that small and slow rising Ca^2+^ events (red arrows) are not accompanied by EAPs. **B)** The detected spikes from A with expanded time bases and ordered by the integral of the EAP. For clarity, Ca^2+^ spikes were truncated (grey dashed line) at the point of a subsequent Ca^2+^ spike. **C)** The correlation between the peak of the EAP integral and the Ca^2+^ spike amplitude for the cell in A had a R^2^ of 0.76. **D)** The mean R^2^ for the correlation between EAP integral and Ca^2+^ spike amplitude was 0.51 ±0.14 (n=5, N=5) and the strength of this relationship depended on the signal-to-noise ratio of the Ca^2+^ recordings. **D)** The Ca^2+^ spikes from A normalised at the 10-90 % risetime of GCaMP6f indicate a

### T-type Ca channels underpin the small amplitude spikes in CSFcNs

Blockade of HVA Ca^2+^ channels reduced the amplitude of Ca^2+^ spikes and their variance, yet smaller Ca^2+^ spikes remained (Fig. 4). Cd^2+^ at 100 µM effectively blocks all HVA Ca^2+^ channels but has negligible effects on low voltage-activated (LVA) T-type Ca^2+^ channels (Huang, 1989). We next tested if T-type Ca^2+^ channels were responsible for the smaller amplitude Ca^2+^ spikes. The selective T-type Ca^2+^ channel blocker (Xiang et al., 2011) ML218 (3 µM) dramatically reduced spontaneous spiking in CSFcNs (Fig. 7A & B, control = 0.058 ±0.08 Hz vs +ML218 = 0 ±0.0017 Hz, median ±IQR, p = 5.6×10^−6^, Wilcoxon signed rank, n=27, N=4). ML218 also decimated spontaneous EAPs in all CSFcNs tested (Fig. 7 C & D, control = 0.48 ±0.13 Hz vs +ML218 = 0.003 ±0.005 Hz, mean ±SD, p = 0.005, n=4, N=4). This finding was surprising as it implied that voltage-activated sodium channels make minimal contribution to spontaneous somatic spiking in CSFcNs. We confirmed this using 1 µm tetrodotoxin (TTx) which had no significant effect on the amplitude (control 0.76 ±1.28 ΔF/F vs +TTx 0.63 ±1.36 ΔF/F, median ±IQR, p = 0.61, Wilcoxon signed rank, n= 37, N=4) nor frequency (control 0.085 ±0.1 Hz vs TTx 0.104 ±0.1 Hz, median ±IQR, p = 0.32, Wilcoxon signed rank, n= 37, N=4) of spontaneous Ca^2+^ spikes in CSFcNs (Fig. 7E-G). Together this data indicates that, at their soma, CSFcNs generate Ca^2+^ spikes rather than the classical Na^+^ spikes.

**Figure 7:**
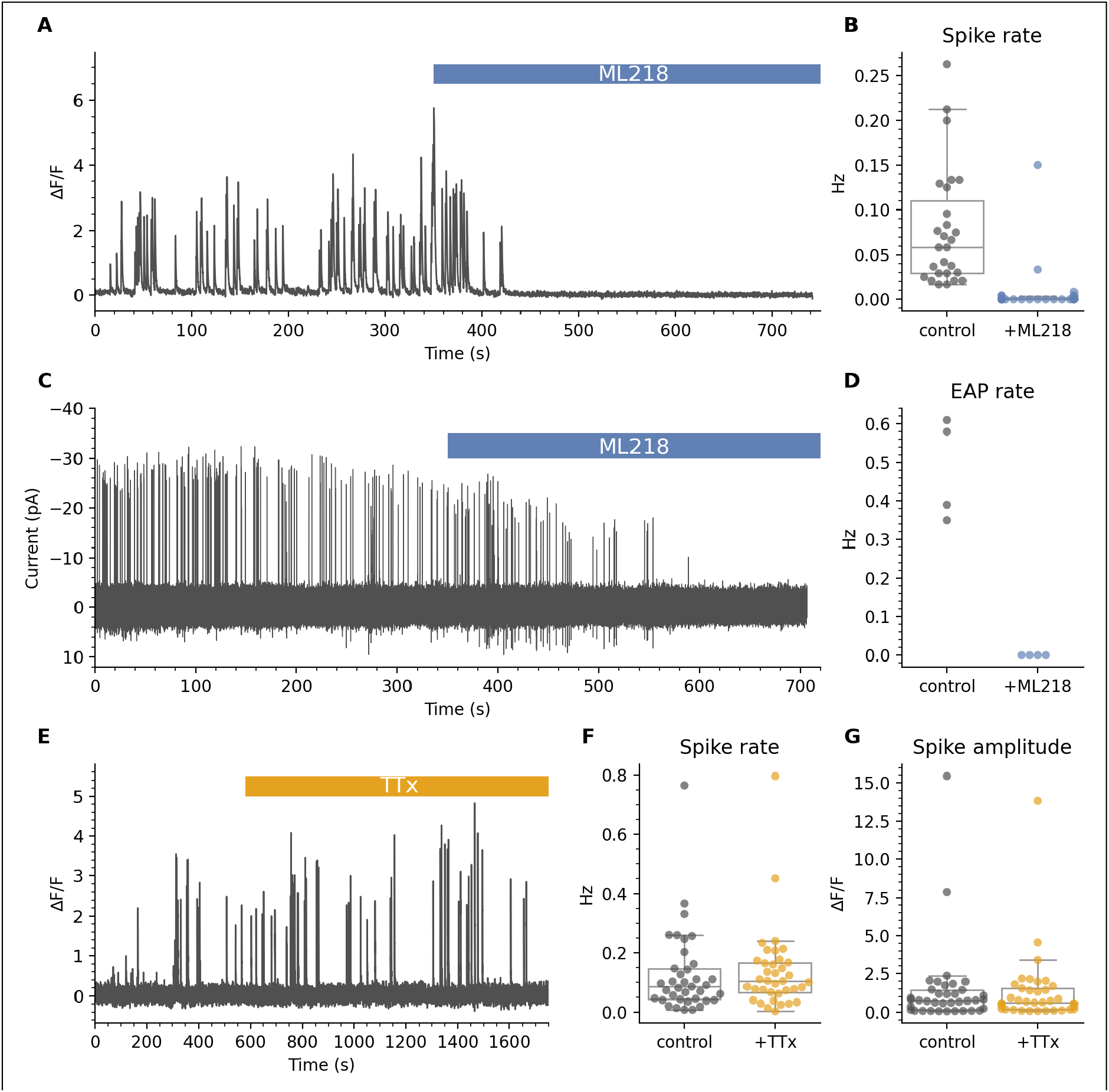
T-Type Ca^2+^ channels are required for spontaneous activity in CSFcNs. **A)** Ca^2+^ recording from a CSFcN before and during application of 3 µM ML218. **B)** ML218 caused a significant reduction in spontaneous firing in CSFcNs, p = 5.6×10^−6^, Wilcoxon signed rank test, n=27, N=4. **C)** EAP recording from a CSFcN before and during application of 3 µM ML218. **D)** ML218 caused a significant reduction in spontaneous EAPs in CSFcNs, p = 0.005, n=4, N=4. **E)** Ca^2+^ recording of a CSFcN before and during application of 1 µM TTx. **F & G)** TTx had no effect on CSFcN spontaneous spike rate or on the amplitude of spikes n = 37, N = 4.

### Spike amplitude as a signal for neurotransmitter activation

We have shown that CSFcNs can generate spikes of different amplitude due to differential recruitment of two types of voltage-activated Ca^2+^ channels. We next tested whether CSFcNs could use spike amplitude as a code to signal activation of different neurotransmitter systems. CSFcNs express nicotinic (Corns et al., 2015) and P2X receptors (Stoeckel et al., 2003; Alfaro-Cervello et al., 2012). We stimulated the same CSFcNs with focal ejection of acetylcholine (1 mM) and ATP (300 µM) which consistently evoked Ca^2+^ spikes with markedly different amplitudes; acetylcholine generated larger spikes (4.63 ΔF/F IQR 2.98) than ATP (0.88 ΔF/F IQR 1.28, n=28, N=3, p = 4.716×10^−6^, Wilcoxon signed rank). Saturating concentrations were used for each agonist (Covernton and Connolly, 2000; Ma et al., 2008) and these experiments were conducted in the presence of a cocktail of synaptic antagonists (20 µM NBQX, 50 µM APV, 10 µM GABAzine, 1 µM strychnine, 1 µM atropine) to prevent any recurrent excitation via non-CSFcNs responding to the agonists. The synaptic blocker cocktail did not affect the amplitude of response suggesting the agonists act directly on CSFcNs and not through excitation of presynaptic neurones (ACh: 4.52 ΔF/F IQR 3.03 vs ACH+synaptic blockers: 4.63 ΔF/F IQR 2.98, p =0.716, ATP: 1.51 ΔF/F IQR 2.16 vs ATP+synaptic blockers: 0.88 ΔF/F IQR 1.28, n = 28, N = 3, p=0.151, Wilcoxon signed rank). Standard aCSF focally ejected in place of the agonists did not evoke activity (data not shown), indicating no contribution from mechanical activation at the pressures used.

This data indicates that CSFcNs can use an amplitude code to signal whether cholinergic or purinergic inputs have been activated, with Ach evoking large amplitude spikes and ATP evoking smaller spikes. We have shown that smaller spikes require T-type Ca^2+^ channels (Fig. 7), whereas larger spikes additionally recruit Cd^2+^ sensitive HVA Ca^2+^ channels (Fig. 4). The evoked ATP responses may therefore be particularly sensitive to block of T-type Ca^2+^ channels. Indeed, application of ML218 caused a large attenuation of ATP responses (Fig. 8 ATP: 0.57 ΔF/F IQR 0.68 vs ATP+ML218: 0.04 ΔF/F IQR 0.22, n=35, N=4) but had minimal effect on acetylcholine evoked responses (Fig. 8 ACh: 2.80 ΔF/F IQR 2.24 vs ACh+ML218: 2.42 ΔF/F IQR 2.18, n=64, N=3, p=0.536, Wilcoxon signed rank).

**Figure 8:**
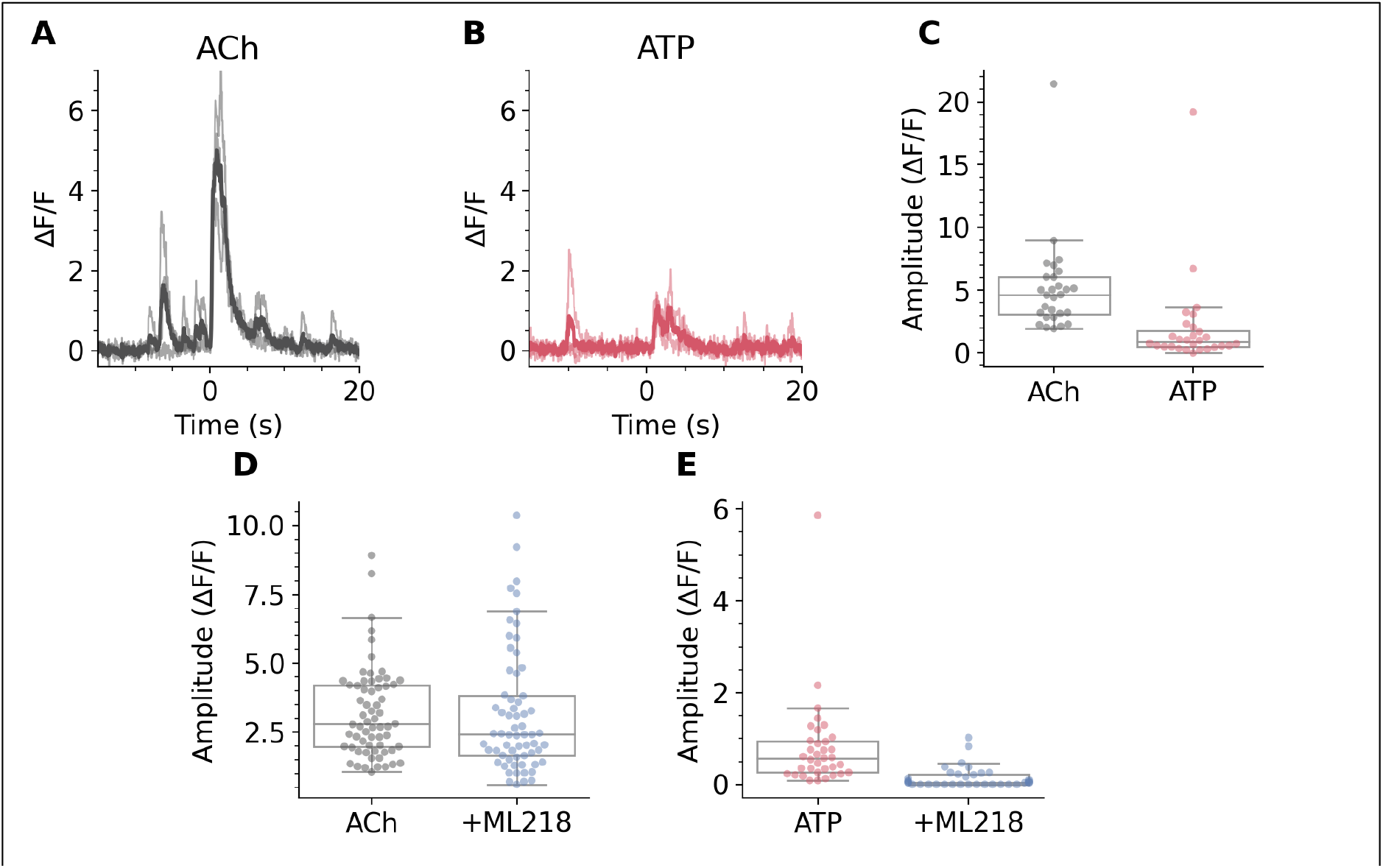
CSFcNs respond to acetylcholine and ATP with different amplitude spikes. **A)** The response of a CSFcN to focal application of Acetylcholine (Ach 1 mM), B) The same CSFcN as in A responding to focal application of ATP (300 µM). For both A & B 3 trial are overlain with the average shown in the dark. Experiment was conducted in the presence of 20 μM NBQX, 50 μM APV, 10 μM GABAzine, 1 μM strychnine and 1 μM atropine. **C)** CSFcNs generated larger spikes to ACh (4.63 ΔF/F IQR 2.98) compared to ATP (0.88 ΔF/F IQR 1.28, n=28, N=3, p = 4.716×10^−6^, Wilcoxon signed rank. **D)** 3 µM ML218 had no effect of Ach evoked responses, 2.80 ΔF/F IQR 2.24 vs 2.42 ΔF/F IQR 2.18, n=64, N=3, p=0.536, Wilcoxon signed rank. **E)** ML218 caused a significant reduction in ATP evoked responses, 0.57 ΔF/F IQR 0.68 vs ATP+ML218: 0.04 ΔF/F IQR 0.22, n=35, N=4.

## Discussion

The spike, or action potential, and its “all-or-none” nature is thought to be a fundamental quantum of neural processing. We show that rather than the typical voltage-activated Na^+^ channels, CSFcNs in the spinal cord of mice use voltage-activated Ca^2+^ channels to generate spikes and that by employing different types of Ca^2+^ channel they can generate graded spikes of variable amplitude. CSFcNs can use these graded spikes to differentially signal which neurotransmitter system is providing their input.

Na^+^ dependent spikes are the ubiquitous mechanism for action potential generation throughout the mammalian brain (Raghavan et al., 2019), whilst Ca^2+^ spikes are typically utilised for dendritic signal integration (Golding et al., 1999; Major et al., 2013; Tran-Van-Minh et al., 2016). In contrast, we find that CSFcNs employ Ca^2+^ spikes in their endbulb and soma. Both our Ca^2+^ imaging and EAP recordings provide localised measurements of activity, recording the Ca^2+^ activity within the soma and endbulb and the somatic membrane currents associated with the action potential respectively (Holt and Koch, 1999; Gold et al., 2006), allowing us to correlate these measurements (Fig. 6). Our data does not preclude a role for Na^+^ spikes in the axons of CSFcNs, which project to the ventral fissure (Stoeckel et al., 2003), but it does suggest that they are absent or at low densities in the soma and endbulb.

In a close parallel, somatic Ca^2+^ spikes are a prominent feature of various other sensory cells including retinal bipolar cells (Baden et al., 2013) and auditory hair cells (Marcotti et al., 2003). Retinal bipolar cells signal with graded analogue signals as well as spikes (Dreosti et al., 2011), both of which are mediated by voltage-activated Ca^2+^ channels (Baden et al., 2011; Hu et al., 2009). Such variations in the amplitudes of Ca^2+^ signals likely support the multivesicular amplitude code used by these synapses (James et al., 2019; Jose Moya-Diaz et al., 2021). Both T-type and L-type Ca^2+^ channels contribute to spiking in retinal bipolar cells (Hu et al., 2009) and in a direct parallel with our findings, T-type channels are required for spontaneous firing while L-type channels influence the shape and duration of Ca^2+^ events (Ma and Pan, 2003). Inner hair cells of the cochlea can signal with graded and spiking responses predominantly through voltage-activated Ca^2+^ channels (Johnson et al., 2011; Marcotti et al., 2003) and spontaneous Ca^2+^-mediated action potentials in hair cells are intrinsically generated and are influenced by both ACh and ATP (Johnson et al., 2011). Since their identification, CSF-CNs have been proposed to be sensory neurons (Kolmer, 1921) and more recent work has demonstrated their functional role in both mechano- and chemo-sensation (Böhm et al., 2016; Fidelin et al., 2015; Hubbard et al., 2016; Jalalvand et al., 2018). It seems then that the adoption of Ca^2+^ spikes to enable variable amplitude Ca^2+^ events is a common feature of neurons across different sensory systems.

Our data show that spontaneous firing at the soma of CSFcNs is dependent on T-type Ca^2+^ channels (Fig. 7). Previous work has implicated spontaneous opening of the large conductance PKD2L1 channel as a means of spike generation in CSFcNs; yet with knockout of PKD2L1, spontaneous firing remained albeit at a lower rate of 0.16 Hz (Orts-Del’Immagine et al., 2017). This rate is identical to what we find when imaging CSFcN activity (the data in Fig. 2G, is not normally distributed but for direct comparison the mean was 0.16 ±0.08 Hz). The PKD family of genes are thought to contribute to mechanosensation (Delmas, 2004), in particular, sensing the shear stress of flow in kidney cells (Nauli et al., 2003) and movement of the CSF in CSFcNs (Sternberg et al., 2018), they may also sense the viscosity of the extracellular matrix (Nigro and Boletta, 2021). It is therefore possible that placement of a patch pipette to record electrical signals may cause inadvertent activation of these channels. Indeed, we find that the placement of a patch pipette against a CSFcN, to form a loose patch recording, elevates the activity of the CSFcN (Fig. 5) and the spike rate we measure in this configuration (0.43 Hz) is almost identical to those made previously with PKD2L1 intact (Orts-Del’Immagine et al., 2016). These observations are therefore consistent with mammalian CSFcNs altering their activity in response to mechanical stimuli as has been described for zebrafish and lamprey (Böhm et al., 2016; Hubbard et al., 2016; Sternberg et al., 2018; Jalalvand et al., 2018).

The functional role of CSFcNs in the mammalian spinal cord is yet to be determined, however they are recipients of numerous axon terminals on their basal pole; including purinergic and cholinergic inputs (Stoeckel et al., 2003; Alfaro-Cervello et al., 2012; Corns et al., 2015). We show that inputs from such purinergic and cholinergic receptors are processed differently by CSFcNs; purinergic inputs are weaker, evoking smaller amplitude T-type dependent Ca^2+^ spikes whereas cholinergic inputs evoke larger spikes that do not depend on T-type channels (Fig 8). A pertinent question is what is the role of these different amplitude spikes in CSFcNs? The apical process within the lumen of the central canal contains vesicles (Jaeger et al., 1983; Alfaro-Cervello et al., 2012) and may release GABA from this site; in lamprey CSFcNs containing somatostatin dense core vesicles appear to release these from their endbulb/soma (Jalalvand et al., 2022). CSFcNs also send an axon to the ventromedial fissure where their synaptic terminals intermingle with axons of the corticospinal tract (Stoeckel et al., 2003). How might the different spike types we describe contribute to transmitter release? Typically, the HVA P/Q and N-type Ca^2+^ channels are coupled to transmitter release in neural cells and N-type channels are present in CSFcNs (Jurčić et al., 2019). However, numerous examples of T-type Ca^2+^ channel activity governing transmitter release exist, including in: neuroendocrine cells in the pituitary (Tomić et al., 1999), adrenal glands (Mlinar et al., 1993), retinal bipolar cells (Pan et al., 2001) and olfactory bulb neurons (Johnston and Delaney, 2010; Fekete et al., 2014). It is therefore possible that the smaller T-type Ca^2+^ spikes that we observe in CSFcNs are able to evoke release of vesicles, perhaps with the HVA channels providing higher vesicle release rates or release of dense-core vesicles. Such dual modes of release have been described in chick auditory hair cells where T-type Ca^2+^ currents regulate rapid vesicle release and L-type Ca^2+^ channels regulate sustained neurotransmitter release (Levic and Dulon, 2012). Furthermore, the N-type Ca^2+^ channel in medullar CSFcNs can be modulated by GABA_B_ receptors (Jurčić et al., 2019), which provides further means for CSFcNs to modulate their spike properties dependent on their synaptic input.

In conclusion, we provide novel evidence that CSFcNs use T-type Ca^2+^ channels to generate spontaneous activity and to respond to low amplitude inputs. We reveal how CSFcNs use graded Ca^2+^ spikes to respond to inputs from different neurotransmitter systems via distinct mechanisms. These observations closely mirror findings within other sensory systems and are consistent with CSFcNs functioning as a multi-modal sensory neuron within the mammalian spinal cord.

## Methods

### Animals

Animal handling and experimentation was carried out according to UK Home Office guidelines and the requirements of the United Kingdom Animals (Scientific Procedures) Act 1986. Mice were housed under a 12:12 h light/dark cycle with free access to food and water. All efforts were made to minimize animal suffering and the number of animals used. Vesicular GABA transporter-IRES-Cre mice (*VGAT*.*Cre*, stock 028862, B6J.129S6(FVB)-Slc32a1<tm2(cre)) were crossed with floxed GCaMP6f mice (*GCaMP6f*.*flox*, stock 028865, B6J.CgGt(ROSA)26Sor<tm95.1 (CAGGCaMP6f)), to generate VGATxGCaMP6f mice. Both mouse lines were originally from Jackson Laboratory (Maine, USA) and maintained in house.

### Immunohistochemistry

3 VGATxGCamP6f mice at P18 and 1 each at P30, P46 and P52 were anaesthetised with a terminal dose of sodium pentobarbital (100mg.kg-^1^, I.P, Euthatal, Merial Animal Health, Dublin) and transcardially perfused, initially with phosphate buffer (PB, 0.1M) to remove blood, and then with paraformaldehyde (PFA, 4% in 0.1M PB, 250ml). Brains and spinal cords were removed and post-fixed overnight in PFA. Spinal cords were serially sectioned (40µm, VT1200 vibrating microtome, Leica Microsystems, Milton Keynes, UK) and stored in PBS at 4°C. Sections were incubated with anti-PKD2L1 (1:500, rabbit, Proteintech) and anti-GFP (1:1000, chicken, Abcam) dissolved in PBS with 0.2% Triton X-100 with 5% donkey serum as a non-specific binding blocker. Sections were washed (x3, PBS 10 minutes) before addition of the Alexa Fluor conjugated secondary antibody (1:1000 in PBS, Thermofisher) at room temperature for 2h. Sections were washed (x2 PBS, 10 minutes) before being mounted on microscope slides and allowed to air dry. Sections were covered using vectashield with DAPI (VectorLabs, cat no. H-1800) and a coverslip was added and sealed using nail varnish.

### Acute slice preparation

VGATxGCamP6f mice (P19-P47, both sexes) were terminally anaesthetised with sodium pentobarbital (as above) and decapitated. The spinal column was dissected and spinal cords were hydraulically extruded via pressure ejection of oxygenated (95% O_2_ : 5% CO_2_) sucrose artificial cerebrospinal fluid (sucrose-aCSF, 30°C, 26mM NaHCO_3_, 2.5mM NaH_2_PO_4_, 3mM KCl, 217mM sucrose, 10mM glucose, 2mM MgSO_4._7H_2_0, 1mM CaCl_2_) through the caudal end of the spinal canal, via a 25ml syringe. Thoracolumbar spinal cord (T5-L3) was embedded in agar (1.5% in sucrose-aCSF) and sectioned transversely using a vibrating microtome (400µm, Integraslice 7550, Campden Instruments, Loughborough, UK), in sucrose-aCSF (30°C). Spinal cord sections were transferred to a submerged incubation chamber containing standard aCSF (124mM NaCl, 26mM NaHCO_3_, 10mM glucose, 3mM KCl, 2.5mM NaH_2_PO_4_, 2mM MgSO_4._7H_2_0, 2mM CaCl_2_, room temp), for ≥1 hour prior to recording. *2-photon Ca*^*2+*^ *imaging:* Spinal cord slices were transferred to the recording chamber, of a custom built 2-photon laser scanning microscope, and perfused with oxygenated aCSF (room temp, 20ml/min), driven by a peristaltic pump. GCaMP6f fluorescence was excited at 910 nm using a pulsed Mai Tai eHP DeepSee Ti:sapphire laser system (SpectraPhysics, Santa Clara, CA, USA). A resonant-galvo mirror assembly (Sutter instruments, UK) scanned the beam through a 16x water-dipping objective (N16XLWD-PF, NA 0.8, Nikon, Tokyo, Japan). Fluorescence was detected using GAasP photomultiplier tubes and appropriate filters and dichroic mirrors. Images were acquired at 30-120Hz, using ScanImage v.5 software (Pologruto et al., 2003). Focal ejection of agonists was achieved using microinjection patch electrodes (3-4µm tip diameter at 2-4 psi) connected to a picospritzer II (Parker, USA) and positioned above the central canal. Agonist were dissolved in aCSF and 12.5 µM Sulforhodamine 101 was included to visually confirm drug delivery.

### Extracellular action potential recordings

Cell attached recordings were made using an axopatch 200B (Molecular Devices, Sunnyvale, CA, USA), signals where digitised using a NI-6356 A-D converter (National Instruments, Austin, TX, USA) controlled by Neuromatic software (Rothman and Silver, 2018), which runs in Igor Pro (Wavemetrics, Lake Oswego, OR, USA). Recordings were made with patch pipettes (3-5 MΩ) containing aCSF and 10 µM alexa-594 (Thermofisher, Waltham, MA, USA) and CSFcNs were targeted under visual guidance using their fluorescence. Light suction was applied to form a seal (10-200 MΩ) and extracellular action potentials (EAPs) were recorded in voltage-clamp mode with the command voltage set to give 0 current. This configuration, 0 current through the pipette and a loose seal, is the optimum for ensuring no depolarisation of the cell being recorded (Perkins, 2006). EAPs are shown inverted so that depolarising phase is rising and hyperpolarising phases are falling.

### Drugs and chemicals

All chemicals and drugs were purchased from Sigma-Aldrich (Gillingham, UK) unless otherwise stated.

#### Quantification and statistical analysis

*Immunohistochemical analysis:* For each section a z-stack was taken with a LSM880 confocal (Zeiss, Oberkochen, Gemany). Using FIJI (Schindelin et al., 2012) CSFcNs were manually counted and were identified in the VGAT-GCaMP6 channel as cells possessing a single bulbous apical process extending into the central canal.

### Image segmentation of CSFcNs

The suite2p pipeline v0.10.1 (Pachitariu et al., 2016) was used to extract raw fluorescence time courses from CSFcNs. Data was first registered using the default options (‘nimg_init’: 200, ‘batch_size’: 200, ‘maxregshift’: 0.1, ‘smooth-sigma’: 1.15). ROI detection was then performed using a spatial scale of 24-48 pixels. The ROI masks corresponding to CSFcNs where visually checked and non-CSFcNs, defined as cells lacking a single bulbous apical process extending into the central canal, were excluded from further analysis. Suite2p and related algorithms rely on sparse asynchronous neural activity to segment cells and we found this approach worked well for the spontaneous spiking data shown in figures 2 and 4-7. However, for the data in figure 8, the focal application of agonists generated synchronous activity across many CSFcNs, we therefore extracted fluorescent time courses for these experiments by manually drawing the cells in FIJI (Schindelin et al., 2012). Extracted fluorescent traces were normalised as ΔF/F using the following equation: F-F^0^/F^0^, where F is the raw florescent trace and F^0^ is the baseline fluorescence which we defined as the 15^th^ percentile of the raw fluorescence.

### Spike detection

Ca^2+^ spikes were detected using the differentiated fluorescence signal as shown in figure 2C. The fluorescence trace was first filtered with a 3^rd^ order Savitzky-Golay filter with a 300 ms window. This signal was then differentiated and point-to-point fluctuations were supressed with a 5 point median filter and then filtered with a 3^rd^ order Savitzky-Golay filter with a 166 ms window. SciPy’s ‘signal.find_peaks’ function was then used to detect spikes with a threshold of 5 times the median-absolute-difference of the differentiated signal. EAPs were detected on the band-pass filtered signal (6-1500 Hz), again using ‘signal.find_peaks’ with the threshold set independently for each cell.

### Spike properties

Ca^2+^ spike amplitudes were measured as the difference between the peak ΔF/F signal occurring in a 333ms window around the spike time and the mean signal in the preceding 100ms (grey arrows in Fig. 2C). The rise times of CSFcN was calculated as the duration to rise from and fall to 5% of the peak amplitude of the average EAP waveform, which corresponds to the depolarising phase of the action potential.

### Statistical analysis

To capture the multi-modality of the amplitude distributions for CSFcNs we used the Jarque-Berra statistic (Fig. 2F) and plot the critical value for a significance level of 0.05 (shaded area) obtained from the Jarque-Bera simulation in Igor Pro. If all data were normally distributed only 5% of points would be outside the shaded area. For graphs of summary metrics individual cells are shown as round markers and box plots represent the median, 25^th^ and 75^th^ percentiles, whiskers extend by 1.5 x interquartile range. All stated summary statistics are either mean ± standard deviation or median ± interquartile range as appropriate and stated in the text. ‘n’ is used to represent a single cell and ‘N’ is used to represent an animal. Where significant differences are reported the statistical test is stated after the p value.

## Acknowledgements

We thank all members of the Johnston and Deuchars labs for useful discussions on this work.

This work was supported by the following grants: SBF002\1033, MR/V003747/1 and WT104818MA

---

**Figure S1.**
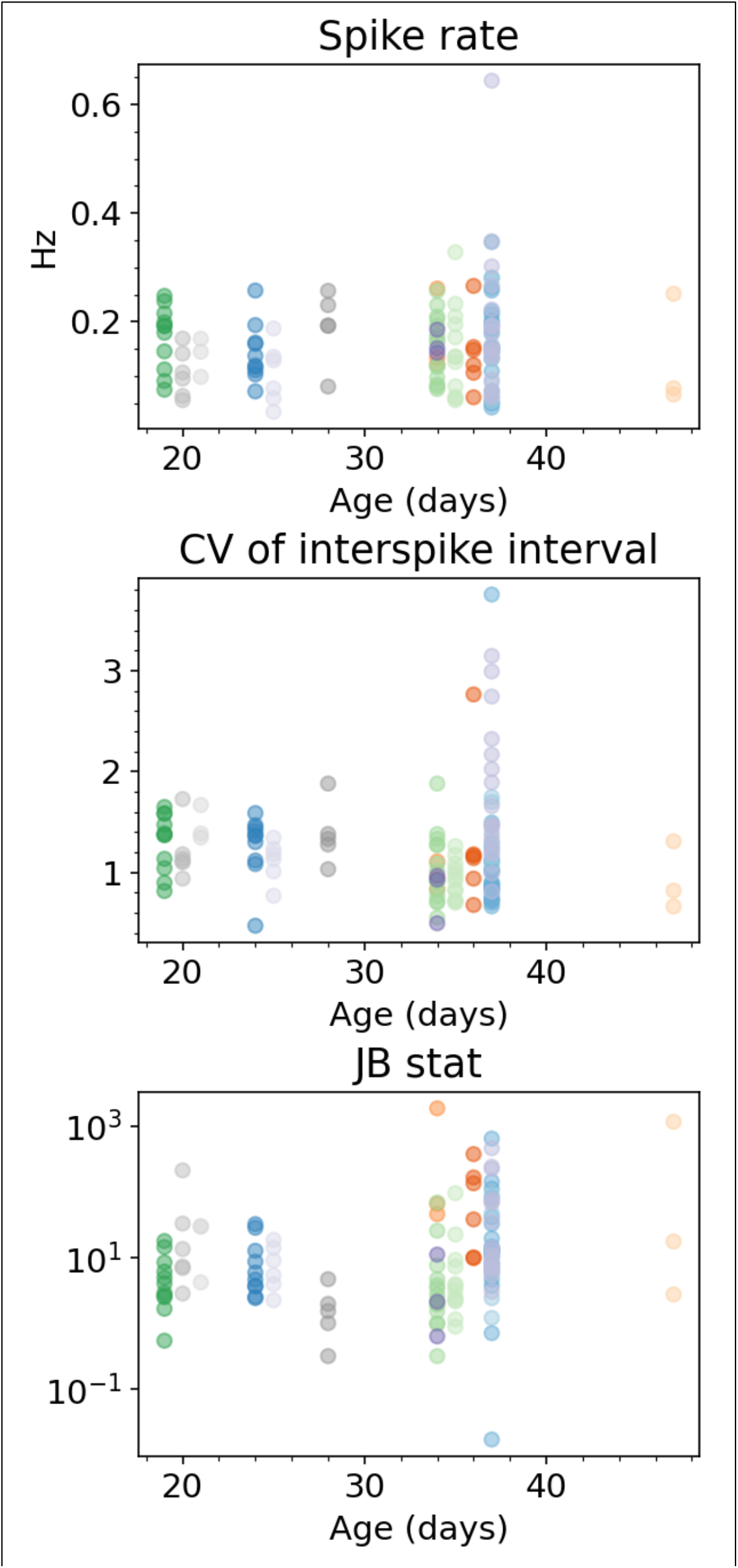
No age dependence of spike metrics. The spike rate, CV of inter-spike-interval and JB statistic were not correlated with the age of animal, p = 0.54, p = 0.21 and p = 0.38 respectively. For each graph different colours represent different animals.

